# Multiplatform Computational Analysis of Mast Cells in Adrenocortical Carcinoma Tumor Microenvironment

**DOI:** 10.1101/2021.02.09.430462

**Authors:** Jordan J. Baechle, David N. Hanna, Konjeti R. Sekhar, Jeffrey C. Rathmell, W. Kimryn Rathmell, Naira Baregamian

## Abstract

**Introduction:** Immunotherapeutic response failure of adrenocortical carcinomas (ACC) highlights a need for novel strategies targeting immune cell populations in the tumor microenvironment (TME) to overcome tumor resistance and enhance therapeutic response. A recent study explored a new link between tumor mast cell (MC) infiltration and improved outcomes in patients with ACC. We further dissect the role of MC in TME of ACC by examining the tumor MC expression signatures and MC activity within TME to provide additional insight into potential novel immunotherapeutic targets.

**Methods:** Using *CIBERSORTx* computational immunogenomic deconvolution algorithm to analyze ACC tumor gene mRNA expression data (TCGA, N=79), we estimated the abundance of tumor immune infiltrating MC, and assessed prognostic potential of MC signaling genes as pro- or anti-tumor signatures, as well as the impact on overall (OS) and disease-free (DFS) survival.

**Results:** We stratified MC signaling genes with survival prognostic values (OS, DFS, p<0.05) into anti-tumor (ALOX5, CCL2, CCL5, CXCL10, HDC, IL16, TNF, TPSAB1, VEGFD) and pro-tumor (CXCL1, CXCL3, CXCL8, IL4, IL13, PTGS3, TNSF4, VEGFD) groups. Anti-tumor MC signature, as the predominant phenotype, was associated with improved OS and DFS.

**Conclusion:** The deconvolution analysis of TCGA data identified MC infiltration in ACC microenvironment as predominantly associated with anti-tumor activity. Future studies stemming from our findings may help define the role of MC in TME and the impact on patient survival in patients with ACC. Modulation of tumor MC infiltration may serve as a potential target for novel synergistic immunotherapies for the treatment and improved survival of patients with ACC.

## INTRODUCTION

Adrenocortical carcinoma (ACC) is among the rarest and most aggressive cancers. Surgical resection remains the mainstay for patients with ACC, but nearly 30% of patients present with unresectable metastasis and resort to medical management^1^. Mitotane therapy, a cytotoxic agent developed in the late 1950s, remains the only approved ACC treatment by the U.S. Food and Drug Administration^2^, and overall survival remains dismal with combination chemotherapy in advanced ACC^3^. Although in recent years immunotherapy has made great strides in many cancer types, the impact of immunotherapy in ACCs has been less encouraging^4,5^, as demonstrated in phase II clinical trial of pembrolizumab efficacy and safety in advanced ACC^4^ and phase IB JAVELIN solid tumor trial (Avelumab) in patients with previously treated metastatic ACC^5^. Response to immunotherapy is dependent on the interaction between tumor cells and tumor microenvironment (TME). The diminished efficacy of immunotherapy directed at key regulators of T cells in ACC was further substantiated by the *Landwehr et al*. study that revealed that the glucocorticoid excess is associated with tumor T cell depletion and unfavorable prognosis^6^. The molecular drivers of immunotherapeutic resistance of ACC are also thought to be at play and include overactivation of the Wnt/ß-catenin pathway or loss of p53 that alter the ability of ACC cells to recruit BAFT3, dendritic cells (DC), and reduced T-cell infiltration^7^.

Tumor mast cell infiltration (TMCI) has recently emerged as a potential prognostic marker of improved outcomes in patients with ACC. *Tian et al*. demonstrated that the tumor mast cell (MC) infiltration is associated with improved outcomes in ACC patients by employing *CIBERSORTx* computational immunogenomic tumor-infiltrating immune cell (TIIC) estimation of the TCGA database and validating the positive prognostic value of TMCI in an independent, single-center cohort of ACC patients^8^. This study provided critical initial insight into the clinical relevance of MC infiltration in TME of ACC, thus necessitating further studies and a deeper understanding of the role of MCs in ACC. In our study, we examined the MC activity within the ACC microenvironment by analyzing the expression of the MC signaling genes using a next-generation *CIBERSORTx* immunogenomic deconvolution analytic platform and explored the MC pro- and anti-tumor effects in the ACC microenvironment. Our preliminary analysis of the ACC large-scale genomic dataset offers a new prism of understanding how MC anti-tumor signaling may potentially be modulated, beyond conventional immunotherapy directed at key regulators of T cells, to optimize immunotherapeutic susceptibility in the ACC microenvironment and improve patient survival in patients with ACC.

## METHODS

### Data Acquisition

We utilized a publicly available, deidentified database with RNA sequencing count table data of Adrenal Cortical Carcinomas (N=92) from The Cancer Genome Atlas (TCGA) Firehose Legacy Cohort^13,14^ through the cBioPortal (https://www.cbioportal.org/) (https://www.cbioportal.org/study?id=60513bc6e4b015b63e9e0b74)^14^. Within the TCGA dataset, only 79 (86%) ACC patients had reported mRNA expression values and were included in the study cohort. This study was exempt from the institutional review board (IRB) approval and did not require patient consent.

### CIBERSORTx Computational Assessment of Prognostic Value of Tumor Mast Cell Infiltration

*CIBERSORTx* immunogenomic deconvolution analytic platform is the next generation of the *CIBERSORT* platform used in the analysis of ACC data by *Tian et al*^*8*^. We employed *CIBERSORTx* to estimate tumor-infiltrating immune subsets in TCGA ACC tumor cohorts. *CIBERSORTx* is a publicly available web-based immunogenomic deconvolution analytic program that uses bulk sample gene mRNA expression data to estimate immune cell abundance and proportional make-up of immune cell subtypes (including B cells, CD4^+^T cells, CD8^+^T cells, dendritic cells, macrophages, natural killer cells, neutrophils, etc.) within TME (https://cibersortx.stanford.edu)^15^.

Tumor mast cell infiltration (TMCI) was analyzed using univariate and multivariate cox regression analysis for its relation to patient overall survival (OS) and disease-free survival (DFS). The predictive prognostic value of TMCI in OS and DFS was determined by a maximally selected log- rank statistic test^16^. All survival and optimal cut-off analyses were performed using the 1.1.383 R statistics software (R Core Team Vienna, Austria).

### Patient Demographics, Tumor Pathology, and Treatment Parameters by Low and High Tumor Mast Cell Infiltration

The American Joint Commission on Cancer (AJCC) Staging Manual, 8^th^ edition, was used to determine TNM classification. Overall survival (OS) was defined as the time from the date of index operation to the date of death. Disease-free survival (DFS) was defined as the time from index operation to the date of documented disease recurrence or death. Categorical variables were presented as frequency and percentages and compared using Chi-square or Fisher’s exact test, as appropriate. Continuous variables were reported as median values with interquartile range (IQR) and compared using the Kruskal–Wallis test. OS and DFS were calculated using the Kaplan–Meier method and compared using the log-rank test. Individual gene expression profiles were analyzed using univariant Cox regression. Significance was set at a *p-*value less than 0.05.

### Immunogenomic Deconvolution of Mast Cell Signaling Gene (MCSG) mRNA Expressions and Tumor-Infiltrating Immune Cells (TIIC) Correlations

All intercorrelations between MC signaling gene (MCSG) mRNA expressions and tumor-infiltrating immune cells (TIIC) were constructed in heatmap format to represent all potential associations using a *CIBERSORTx* platform. Correlations were calculated using Spearman rank correlation coefficient and evaluated using the Pearson correlation formula. Correlational significance was set at a *p-*value less than 0.01 and correlations with significant correlations are represented by heatmap shading. Statistically insignificant heat map correlations (p≥0.01) were void of shading.

### Mast Cell Signaling Gene (MCSG) mRNA Expression Signature Calculation and Proportional Predominance

Cumulative anti- and pro-tumor signatures were normalized to positive values (0-5) and weighted against their sum to quantity the proportion of anti- and pro- mRNA expression signatures in each patient. Expression predominance was determined by proportional expression greater than 50%. OS and DFS were compared according to signature predominance using Kaplan-Meier and Log-rank test.

## RESULTS

### Tumor Mast Cell Infiltration (TMCI) in ACC

Tumor mast cell infiltration (TMCI) was associated with prolonged DFS (Hazard Ratio (HR) 0.02, 95% Confidence Interval (95% CI) 0.00 – 0.64 p=0.026). TMCI was not associated with improved OS (HR 0.27, 95% CI 0.02 – 4.99, p=0.378). Maximally selected log-rank statistic test identified MC infiltration >0.15au (arbitrary units) to be increasingly associated with prolonged DFS and a TMCI value of 0.21 to carry the maximum DFS prognostic value (**Figure 2A, 2B**). Patients with TMCI >0.21 were deemed “high TMCI” group and patients with TMCI <0.21 were considered “low TMCI” group. Low TMCI was associated with shortened DFS (HR 5.51, 95% CI 1.32 – 22.97, p=0.019). No dichotomous threshold of mast cell inflammation was found to be a predictor of OS and OS log-rank comparison using optimal DFS threshold (p=0.17) (**Figure 2C**). DFS was shortened for ACC patients with suboptimal TMCI (5-year DFS, 85.7 vs. 37.1%, p=0.005, **Figure 2D)**.

**Figure 1.**
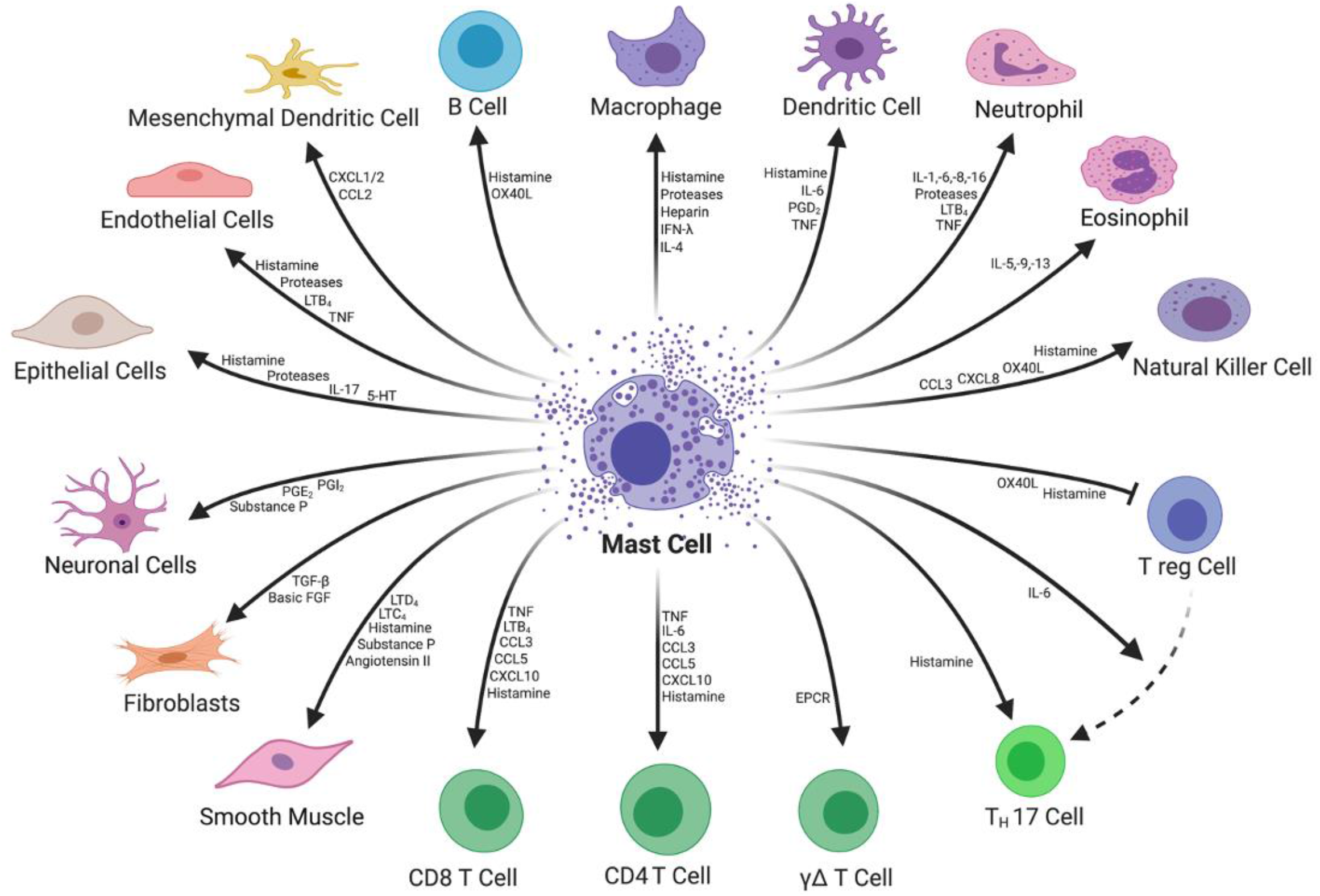
Mast Cell Signaling and Target Cell Types.

**Figure 2.**
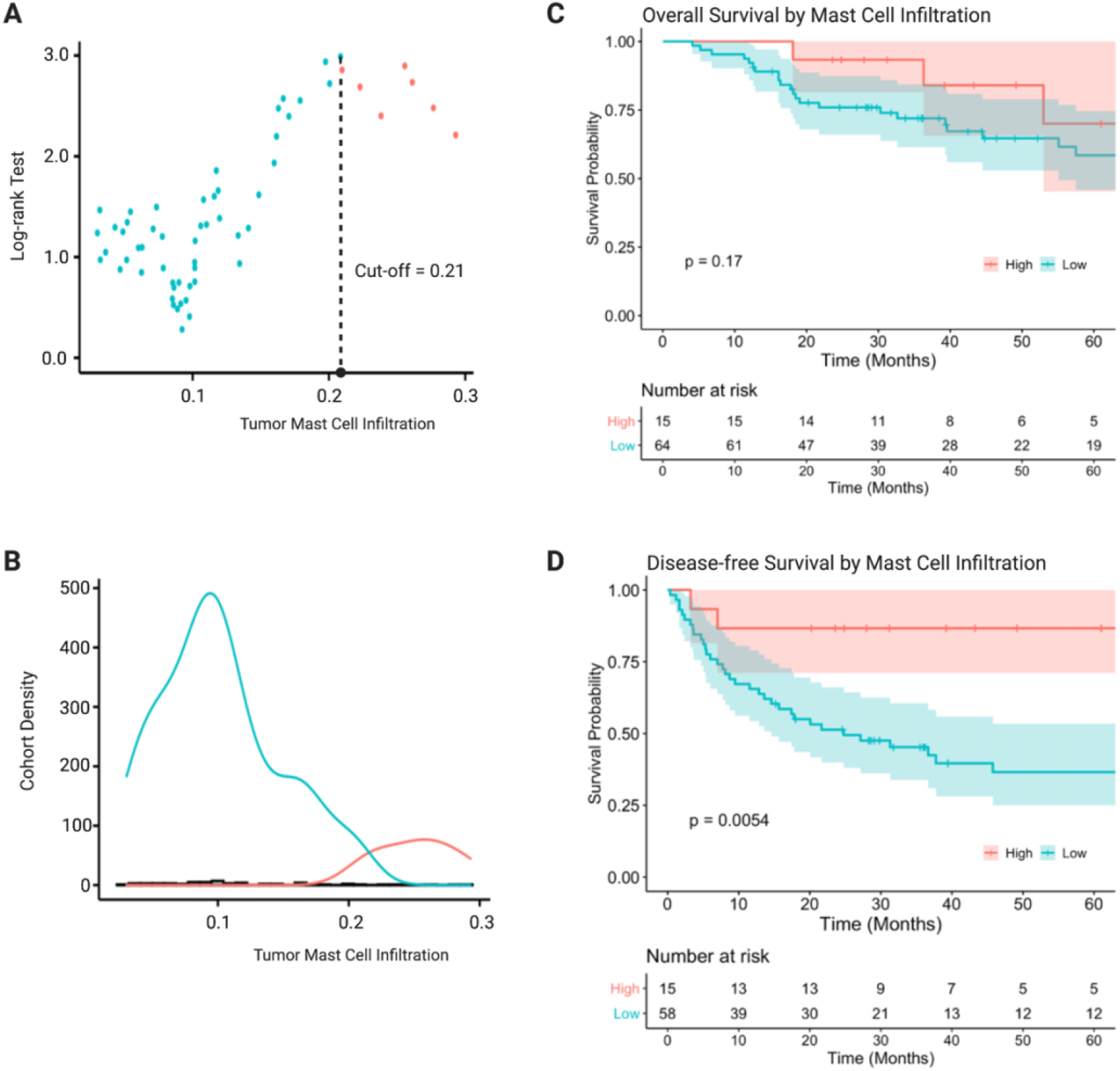
Prognostic Value of Tumor Mast Cell Infiltration in Adrenocortical Carcinoma. **A**. Optimal value of tumor-infiltrating mast cell prognostic for DFS determined by a maximally selected log-rank statistic test. **B**. Distribution of suboptimal (low) and optimal (high) tumor mast cell infiltration within the study cohort. **C**. OS stratified by optimal and suboptimal mast cell infiltration. **D**. DFS stratified by optimal and suboptimal mast cell infiltration.

### Anti-tumor MCSG and Tumor-Infiltrating Immune Cell (TIIC) Profile Correlations

A comprehensive list of MCSG and their survival impact were analyzed using a univariate Cox regression survival analysis and were summarized in **Supplemental Table 2**. MCSG with significant positive prognostic value (Hazard Ratio (HR) < 1.00, p<0.05) in OS or DFS were deemed anti-tumor MCSG. The anti-tumor MCSG included ALOX5, CCL2, CCL5, CXCL10, HDC, IL16, TNF, and VEGFD (**Figure 3A, 3B**). Cumulative anti-tumor MCSG signature expression was associated with prolonged DFS (HR 0.893, 95% CI 0.84 – 0.95, p<0.001) only. The role of anti-tumor MCSG are summarized in **Figure 3C**. There were several significant mRNA expression intercorrelations between all anti-tumor MCSGs (r>0.3) with the exception of VEGFD. VEGFD expression was positively correlated with histidine decarboxylase (HDC) expression (r=0.32) (**Figure 3D**). Expression and TIIC correlations of pro-tumor MCSG revealed anti-tumor MCSG expression to be positively correlated with the infiltration of CD8^+^ T cells (r=0.54), T(reg) cells (r=0.53), M1 (r=0.54) and M2 (r=0.63) macrophage, and total macrophage and monocyte (r=0.64) (p-values <0.01) (**Figure 3E**).

**Figure 3.**
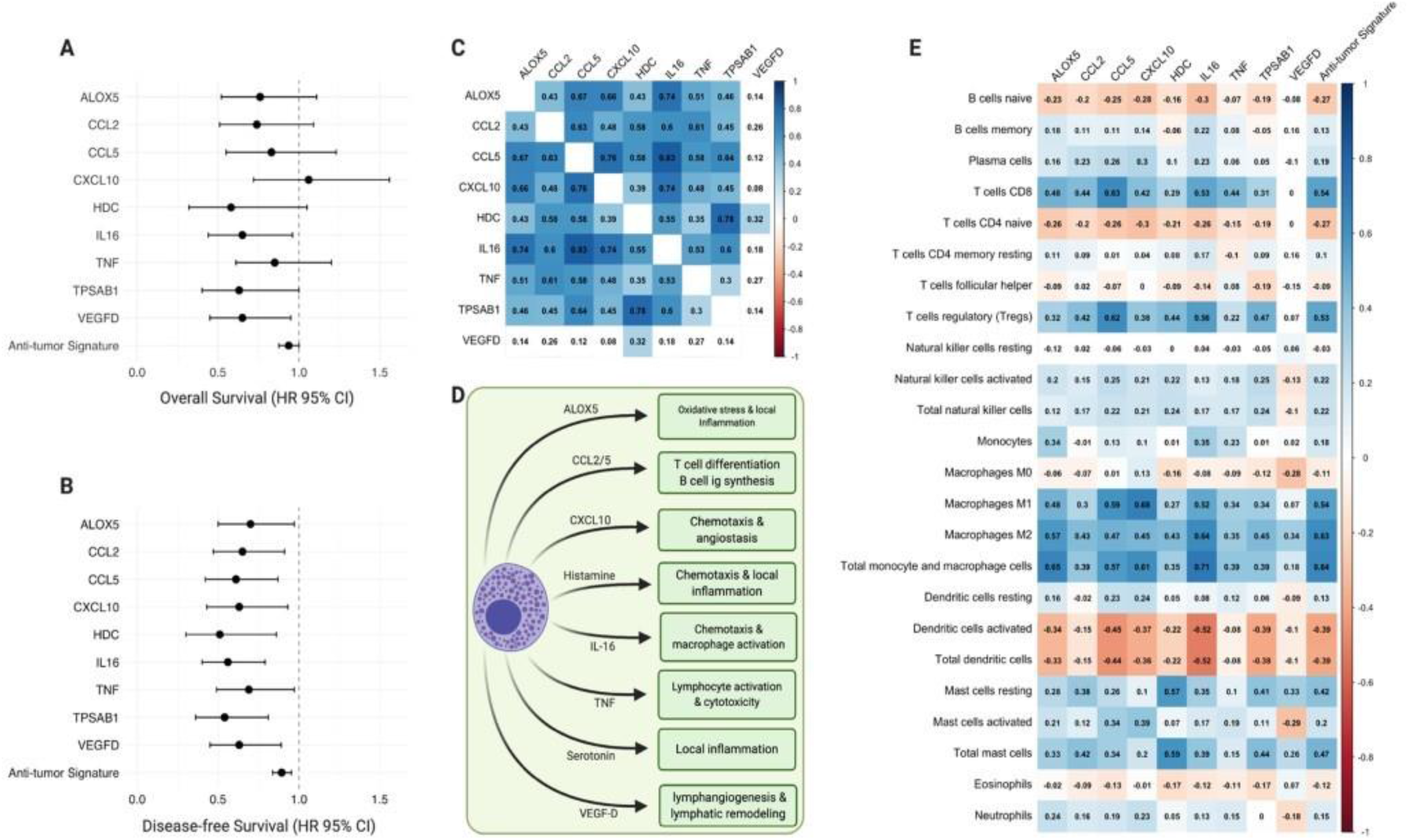
Anti-tumor Mast Cell Signaling Genes (MCSG) in Adrenocortical Carcinoma. **A, B**. Significant prognostic indicators of MCSG with significant associations for overall or disease-free survival (OS or DFS, HR <1.00, p<0.05). **C**. Heatmap of anti-tumor mast cell signaling gene expression showing Spearman rank correlation coefficients with significant correlations (p<0.01) shaded. **D**. Diagram of anti-tumor mast cell activity and anti-tumor mast cell gene signaling roles. **E**. Heatmap of individual gene expression contributing to anti-tumor mast cell signaling signature with tumor-infiltrating immune cell (TIIC) subtypes. Significant correlations (p<0.01) shaded.

### Pro-tumor MCSG and TIIC Profile Correlations

MCSG with significant negative prognostic value (Hazard Ratio (HR) > 1.00, p<0.05) in overall (OS) or disease-free survival (DFS) were deemed pro-tumor MCSGs. These genes included CXCLl, CXCL8, IL4, IL13, PTGS2, TNFSF4, and VEGFC. Cumulative pro-tumor MCSG signature expression was significantly associated with shortened OS and DFS (OS HR 1.18, 95% CI 1.09 – 1.29, p<0.001; DFS HR 1.14, 95% CI 1.05 – 1.23, p<0.001) (**Figure 4A, 4B**). The role of pro-tumor MCSGs are summarized in **Figure 4C**. Strong intercorrelations were found between CXCL1, CXCL3, CXCL8, and PTGS2 mRNA expression (r>0.55) (**Figure 4D**). Pro-tumor MCSG expression was positively correlated with the infiltration of M1 macrophage (r=0.53), activated mast cells (r=0.52) and neutrophil (r=0.53) infiltration (**Figure 4E**) (p-values <0.01).

**Figure 4.**
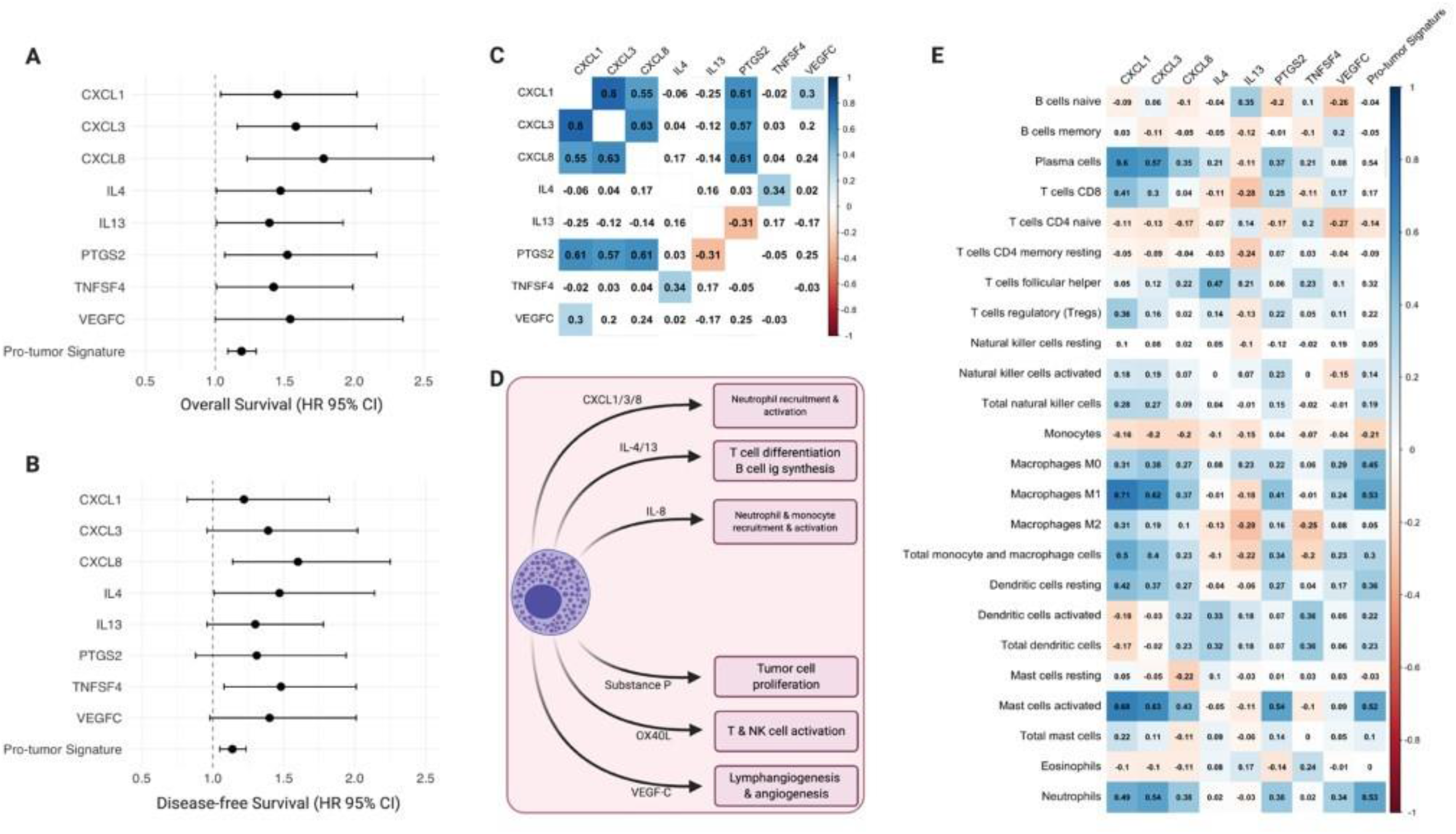
Pro-tumor Mast Cell Signaling Genes (MCSG) in Adrenocortical Carcinoma. **A, B**. Significant prognostic indicators of MCSG with significant associations for overall and disease-free survival (OS or DFS, HR >1.00, p<0.05). **C**. Heatmap of pro-tumor mast cell signaling gene expression showing Spearman rank correlation coefficients with significant correlations (p<0.01) shaded. **D**. Diagram of pro-tumor mast cell activity and pro-tumor mast cell gene signaling roles **E**. Heatmap of individual gene expression contributing to pro-tumor mast cell signaling signature with tumor-infiltrating immune cell (TIIC) subtypes. Significant correlations (p<0.01) shaded.

### Anti-tumor MCSG Expression Signature Predicts Survival Outcomes

Of the entire cohort, 72% (n=57) were identified to predominantly express anti-tumor gene signature and 28% (n=22) were pro-tumor signature predominant (**Figure 5A**). Anti-tumor mast cell signaling predominance among the cohort further suggests that although mast cells may play a heterogenous role in ACC TME, they may be considered a favorable prognostic indicator with a beneficial overall impact on the TME. Anti-tumor and pro-tumor signature predominant groups were made up of similar portions of CS- and nonCS-ACC patients (54.5 vs. 36.8%, p=0.24). Anti-tumor predominant MCSG expression was associated with prolonged 5-year overall (73.0 vs. 28.3%, p<0.001) and DFS (53.9 vs. 9.7%, p<0.001) survival compared to ACC patients with pro-tumor predominant MCSG expression (**Figure 5A, B, C**).

**Figure 5.**
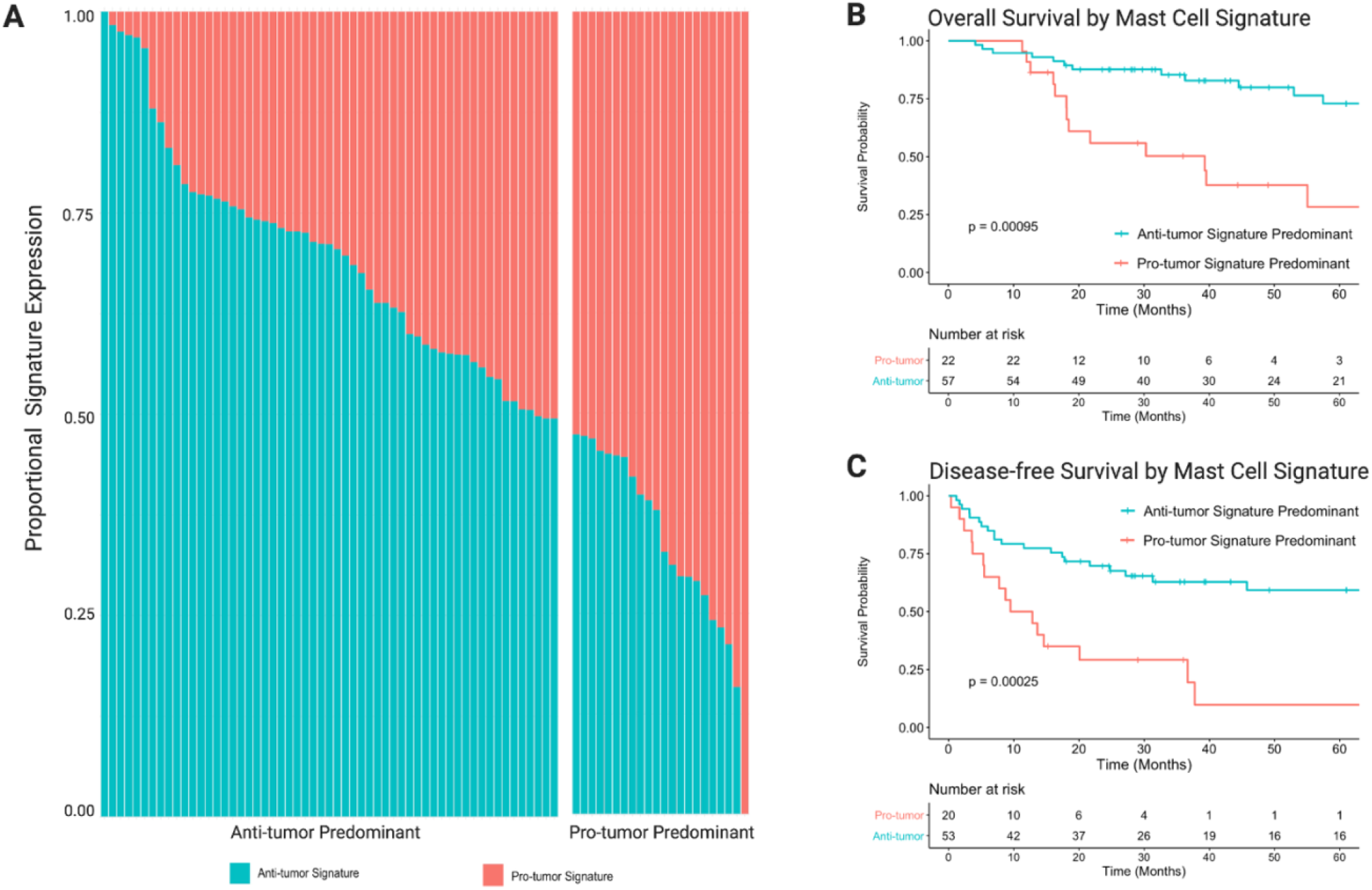
Mast Cell Signaling Gene mRNA Expression Signature Predominance. **A**. Proportional mRNA expression of anti- and pro-tumor MCSG signatures in each patient. **B**. Overall survival comparing anti- and pro-tumor MCSG signature expression predominance **C**. Disease-free survival comparing anti- and pro-tumor MCSG signature expression predominance.

### MCSG Expression Profile of Excess Cortisol Secreting ACC

Among the anti-tumor MCSGs associated with favorable prognosis, the mRNA expression of CCL5 (p=0.008), HDC (p<0.001), IL16 (p=0.002), and TPSAB1 (p<0.001) were downregulated in CS- ACCs compared to nonCS-ACCs (**Supplemental Figure 1A**). Furthermore, the anti-tumor expression signature was decreased in CS-ACC relative to nonCS-ACC patients (p=0.006). CS- and nonCS-ACCs demonstrated similar expression of the pro-tumor MCSGs and the pro-tumor cumulative signature (**Supplemental Figure 1B**).

### Patient Demographic, Tumor Pathology, and Treatment Parameters of ACC: Low vs High TMCI

Clinical and treatment parameters were stratified according to the low vs high TMCI distribution in ACC tumors that may confound DFS and/or OS associations (**Supplemental Table 1**). Among the 79 patients included in the cohort, 64 (81%) had low TMCI and 15 (19%) had high TMCI. The groups were similar in age at diagnosis (p=0.871), sex (p=0.718) tumor stage T (p=0.622), nodal status N (p=0.791), metastasis M (p=1) and clinical stage (p=0.622).The high TMCI group had a greater portion of patients with an unreported race (33.0 vs 9.4%, p=0.014). Low and high TMCI groups demonstrated similar fractions of genome alteration (p=0.276), mutation count (p=0.222), mitotic rate (p=0.556), tumor necrosis (p=1), Weiss Score^17^ (p=0.545), and rates of vascular invasion (p=0.849). Both groups reported similar resection margins (p=0.866) and underwent similar rates of unspecified neoadjuvant (p=1) and adjuvant (p=0.662) therapy, as well as radiation (p=0.866) and mitotane therapy (p=0.231) throughout their clinical course.

Significant differences between the low and high TMCI groups were observed with the type of excess adrenal steroid oversecretion (p=0.047), with higher rates of isolated cortisol-secreting ACC (CS-ACC) among the low TMCI group (25.4 vs 0.0%, p<0.001), higher rates of ACC recurrence following initial treatment (55.9 vs 14.3%, p=0.012). On multivariable Cox regression analysis controlling for cortisol-secretion, low TMCI was associated with shortened DFS (HR 4.37, 95% CI – 18.60, p=0.046). Thus, the poor DFS prognosis related to low TMCI after controlling for cortisol-secretion and despite similar patient demographics, tumor pathology, and treatment protocols commonly associated with survival (patient age, cancer stage, Weiss Score, neoadjuvant/adjuvant therapy) is suggestive of a possible influential role of mast cell activity in ACC TME that may underlie disease biology and patient prognosis.

## DISCUSSION

In this study, we utilized multiplatform computational immunogenomic deconvolution and bulk mRNA expression analysis of TCGA ACC data from cBioPortal to gain a deeper understanding of the role of MC tumor infiltration and signaling activity within ACC TME. Our findings are multifaceted in that they support previous research characterizing tumor-infiltrating MC abundance as a favorable prognostic factor in ACC^8^. Our analyses dive deeper to investigate potential confounders underlying the previously identified favorable prognostic value of MC in ACC TME, and we outline prognostic pro- and anti-tumorigenic MC signaling genes to lay a foundation for future mechanistic studies to investigate and exploit potential MC anti-tumor activity in ACC tumors.

The demographic, clinical, pathologic, and treatment parameter comparisons between ACC patients harboring low and high TMCI, excess cortical steroid secretion emerged as the only factor identified to be significantly different between groups with isolated excess cortisol secretion significantly associated with low TMCI. This finding implied an influential role of glucocorticoid-induced MC immunomodulation, therefore, we performed additional analysis to compare the anti- and pro-tumorigenic MC gene expression levels by cortisol secretion and found several anti-tumor genes (CCL5, HDC, IL16, TPSAB1) to be downregulated in CS-ACCs compared to nonCS-ACCs, as was the cumulative anti-tumor MCSG signature. Excess cortisol secretion is known for both immunosuppression in ACC TME^18^ and as an independent risk factor for poor prognosis in ACC patients^19^. The DFS prognostic value of TMCI, however, maintained statistical significance after controlling for excess cortisol secretion on multivariant analysis, while excess cortisol secretion did not. Together, these correlations indicate the potential for cortisol-induced suppression of anti-tumor MC signaling (among other interactions), which may, in part, contribute to diminished MC anti-tumor activity in ACC TME and the poor prognosis in patients with CS-ACC.

Previous studies have demonstrated a wide range of variability in the prognostic value of MC infiltration and activity according to the type, grade, or stage of the tumor, and the prognostic role MC has across cancer types^9-12,20,21^. MC appears to play a pro-tumorigenic role and an anti-tumorigenic role in various cancers, and to some extent, can be non-contributory in certain tumors. This collage of evidence suggests a heterogeneous role of MC in the immune landscape of tumors that may vary on a spectrum across different tissue and cancer types. The variation may also be influenced by differential MC signaling activity and gene expression, as well as the susceptibility of specific tumor types to inflammation.

Our study assessed the prognostic value of the mRNA expression of genes attributable to MC signaling to characterize the dichotomous pro-and anti-tumorigenic MC activity. The anti-tumor MCSGs associated with improved prognosis were found to be predominantly involved in promoting local inflammation (ALOX5/leukotriene B_4_ HDC/histamine, TPSAB1/serotonin), as well as lymphocyte (CCL2, CCL5, CXCL10, TNF) and phagocyte (IL-16) chemotaxis and activation. VEGF is well known for its angio- and lymphogenic properties in TME, however, the role of differentially expressed VEGF subunits (-A, -B, -C, and -D) remains less understood. In this study, VEGF-D expression was associated with improved prognosis and VEGF-C with poor prognosis. A similar phenomenon has been observed in non-small cell lung cancer and attributed to the increased lymphogenic properties of VEGF-C over VEGF-D^22^. While the VEGF-C expression is considered a risk factor for the lymphatic spread in several cancers, VEGF-D remains less characterized clinically but has demonstrated decreased levels to be associated with metastatic disease, including thyroid tumors^23^, suggesting that VEGF-D may serve as a competitive agonist at the VEGF receptor. Reduced levels of VEGF-D may allow the increased activity of VEGF-A and VEGF-C.

The pro-tumor MCSG associated with poor prognosis was found to be mostly involved in the promotion of neutrophil recruitment and activation (CXCL1, CXCL3, CXCL8, IL-8), and the expression of the pro-tumor signature was strongly correlated with neutrophil tumor infiltration. Substance P (PTGS2) mRNA expression was also associated with shorter OS. Substance P is released upon mast cell degranulation, can induce tumor cell mitogenicity (proliferation, invasion, metastasis) through activation of the neurokinin (NK)-1 receptor system, while NK-1 antagonists negate such effects^25^.

The anti- and pro-tumor MCSG signatures appeared to inversely impact prognosis (OS and DFS) to a similar degree, however, the study cohort was shown to predominantly express the anti- tumor signature over the pro-tumor MCSG signature and anti-tumor predominance was significantly associated with improved prognosis. These findings support the favorable prognostic value of MC infiltration in ACC patients recently identified by Tian and colleagues at Fudan University in Shanghai despite the heterogenous role of MC in TME. Furthermore, our study provides insight into potential contributory factors that may underly the dichotomous role of MC observed across various cancer types. Of the pro- and anti-tumor ACC MCSGs identified in this study, promotion of local inflammation and immune cell recruitment through ALOX5/leukotriene B_4_, HDC/histamine, and TPSAB1/serotonin may be a potential avenue for supplementing MC signaling in ACC TME.

This study is not without limitations. Primarily, this is a retrospective study design examining large-scale sequenced genetic data of ACC tumors using a computational deconvolution approach to ascertain tumor immune cell infiltration within TME and inherently is limited from determining causal relationships. Although we feel that the study’s use of the TCGA database strengthened this study by providing a sufficiently robust database of clinical and genetic parameters to derive meaningful associations regarding the TME of these ultra-rare tumors, the collaborative is limited to large, academic referral centers which may lead to selection bias towards more advanced stage disease with over-representation of metastatic disease. This may skew our findings and impact conclusions in such a way that it can limit the generalizability of our conclusions to more advanced, treatment-resistant ACCs. Furthermore, the collaborative nature of the TCGA database also limits the granularity of clinical data available for certain parameters relevant to clinicians. For example, the TCGA database only reports on ACC hormone hypersecretion (nonfunctional, cortisol, aldosterone, androgen, etc.) and does not include the diagnostic test use or lab values. Similarly, the site of metastasis is unreported in this database. Furthermore, the treatment of patients with ACC is very heterogeneous with variations in surgical technique, radiation therapy (adjuvant/neoadjuvant), mitotane regimen (including dose, frequency, therapeutic level). Our survival analysis was also limited to time from index operation and may not fully grasp the prognosis from date of diagnosis for patients receiving preoperative treatment or otherwise delayed surgical resection. Altogether, such limitations, inherent to large databases of rare tumors, hinder our ability to further characterize and investigate these and other clinical treatment factors that may impact OS and DFS and influence MC infiltration and activity.

Moreover, it is challenging to clearly define the role of individual immune cell types due to overlap in function, gene expression, and immune interplay, and this study is no exception. This study does not include single-cell analysis or special profiling transcriptomics and the expression of genes analyzed in this study are likely not exclusive to MCs. Instead, this study aims to identify the potential activity of (but not limited to) MCs that may underly their prognostic value in ACC TME consistent across multiple methods of immunogenomic and immunohistochemistry quantification^9^. Collectively, our findings pave a way for future studies focusing on uncovering the underlying mechanisms that may influence the MC infiltration, and expression and immunomodulatory activity of MCSG in the ACC TME.

In conclusion, our study provides interesting new insight into the nuanced role of MC infiltration in ACC TME and identifies potential areas worthy of further investigation in independent cohorts. The augmentation or supplementation of anti-tumor MC signaling may be an efficient means of optimizing TME immune response and enhancing immunotherapeutic response in patients with ACC.

## SUPPLEMENTAL TABLES & FIGURES

**Supplemental Table 1.** Patient Demographic, Tumor Pathology, and Treatment Parameters of Adrenocortical Carcinoma. IQR = interquartile range, Age = age at time of index operation, Unknown = unreported data, HPF = high-power fields, RX = indeterminate margins. *Mitotane and *radiation therapy at any point in management.

|  |  | Low Mast Cells<br>N=64 | High Mast Cells<br>N=15 | p-value |
| --- | --- | --- | --- | --- |
| **Age (years) -Median [IQR]** |  | 48.0 [35.5;60.0] | 50.0 [33.5;53.0] | 0.871 |
| **Sex** | Female | 40 (62.5%) | 8 (53.3%) | 0.718 |
|  | Male | 24 (37.5%) | 7 (46.7%) |  |
| **Race** | White | 57 (89.1%) | 9 (60.0%) | 0.014 |
|  | Asian | 1 (1.56%) | 0 (0.00%) |  |
|  | Black | 0 (0.00%) | 1 (6.67%) |  |
|  | Unknown | 6 (9.38%) | 5 (33.3%) |  |
| **Clinical Stage** | Stage I | 7 (10.9%) | 2 (13.3%) | 0.622 |
|  | Stage II | 28 (43.8%) | 9 (60.0%) |  |
|  | Stage III | 15 (23.4%) | 1 (6.67%) |  |
|  | Stage IV | 12 (18.8%) | 3 (20.0%) |  |
|  | Unknown | 2 (3.12%) | 0 (0.00%) |  |
| **T Stage** | T1 | 7 (10.9%) | 2 (13.3%) | 0.821 |
|  | T2 | 32 (50.0%) | 10 (66.7%) |  |
|  | T3 | 7 (10.9%) | 1 (6.67%) |  |
|  | T4 | 16 (25.0%) | 2 (13.3%) |  |
|  | Unknown | 2 (3.12%) | 0 (0.00%) |  |
| **N Stage** | N0 | 54 (84.4%) | 14 (93.3%) | 0.791 |
|  | N1 | 8 (12.5%) | 1 (6.67%) |  |
|  | Unknown | 8 (12.5%) | 0 (0.00%) |  |
| **M Stage** | M0 | 50 (78.1%) | 12 (80.0%) | 1.00 |
|  | M1 | 12 (18.8%) | 3 (20.0%) |  |
|  | Unknown | 2 (3.12%) | 0 (0.00%) |  |
| **Excess Hormone Secretion** | Cortisol | 16 (25.4%) | 0 (0.00%) | 0.047 |
|  | Androgen + Cortisol | 13 (20.6%) | 3 (20.0%) |  |
|  | Androgen | 8 (12.7%) | 0 (0.00%) |  |
|  | Estrogen | 1 (1.59%) | 1 (6.67%) |  |
|  | Mineralocorticoids | 2 (3.17%) | 1 (6.67%) |  |
|  | Mineralocorticoid + Cortisol | 1 (1.59%) | 0 (0.00%) |  |
|  | None | 21 (33.3%) | 9 (60.0%) |  |
| **Excess Cortisol (including mixed) Secretion** | No | 34 (53.1%) | 12 (80%) | 0.108 |
|  | Yes | 30 (46.9%) | 3 (20.0%) |  |
| **Laterality** | Left | 38 (59.4%) | 7 (46.7%) | 0.545 |
|  | Right | 26 (40.6%) | 8 (53.3%) |  |
| **Fraction of Genome Altered -Median [IQR]** |  | 0.60 [0.38;0.91] | 0.42 [0.25;0.83] | 0.276 |
| **Mutation Count -Median [IQR]** |  | 87.0 [72.5;118] | 81.5 [57.5;92.8] | 0.222 |
| **Mitoses Count -Median [IQR]** |  | 7.00 [5.00;23.0] | 7.00 [4.00;15.0] | 0.556 |
| **Mitotic Rate** | < 5/50 HPF | 24 (37.5%) | 5 (33.3%) | 0.777 |
|  | > 5/50 HPF | 32 (50.0%) | 9 (60.0%) |  |
|  | Unknown | 8 (12.5%) | 1 (6.8%) |  |
| **Tumor Necrosis** | Absent | 14 (21.9%) | 3 (20.0%) | 1.00 |
|  | Present | 46 (71.9%) | 11 (73.3%) |  |
| **Weiss Score -Median [IQR]** |  | 5.50 [3.75;7.25] | 7.00 [5.00;7.00] | 0.545 |
| **Venous Invasion** | Absent | 31 (48.4%) | 8 (53.3%) | 0.849 |
|  | Present | 26 (40.6%) | 5 (33.3%) |  |
|  | Unknown | 7 (10.9%) | 2 (13.3%) |  |
| **Neoadjuvant Therapy** | No | 63 (98.4%) | 1 (1.56%) | 1.000 |
|  | Yes | 1 (1.56%) | 0 (0.00%) |  |
| **Resection Margin** | R0 | 43 (67.2%) | 12 (80.0%) | 0.866 |
|  | R1 | 5 (7.81%) | 1 (6.67%) |  |
|  | R2 | 7 (10.9%) | 2 (13.3%) |  |
|  | RX | 6 (9.38%) | 0 (0.00%) |  |
|  | Unknown | 3 (4.69%) | 0 (0.00%) |  |
| **Adjuvant Therapy** | No | 20 (31.2%) | 5 (33.3%) | 0.662 |
|  | Yes | 42 (65.6%) | 9 (60.0%) |  |
|  | Unknown | 2 (3.12%) | 1 (6.67%) |  |
| **Mitotane Therapy\*** | No | 21 (32.8%) | 5 (33.3%) | 0.231 |
|  | Yes | 41 (64.1%) | 8 (53.3%) |  |
|  | Unknown | 2 (3.12%) | 2 (13.3%) |  |
| **Radiation Therapy\*** | No | 47 (73.4%) | 12 (80.0%) | 0.866 |
|  | Yes | 14 (21.9%) | 1 (6.67%) |  |
|  | Unknown | 3 (4.69%) | 2 (13.3%) |  |
| **New Neoplasm Following Initial Treatment** | No | 26 (44.1%) | 12 (85.7%) | 0.012 |
|  | Yes | 33 (55.9%) | 2 (14.3%) |  |

**Supplemental Table 2.** Survival Impact of Mast Cell Signaling Gene Expression in Adrenocortical Carcinoma. OS and DFS; Bold = p<0.05.

| Mast Cell Signal | Primary Gene | Overall Survival<br>HR (95% CI) | P value | Disease-free Survival<br>HR (95% CI) | P value |
| --- | --- | --- | --- | --- | --- |
| 5-HT | TPH1 | 0.99 [0.69;1.41] | 0.938 | 0.99 [0.71;1.38] | 0.938 |
| Angiotensin II | ACE | 0.90 [0.62;1.31] | 0.580 | 0.91 [0.66;1.26] | 0.575 |
| Carboxypeptidase | CPE | 1.02 [0.71;1.45] | 0.928 | 0.76 [0.56;1.04] | 0.092 |
| **CCL2** | **CCL2** | 0.74 [0.51;1.09] | 0.130 | **0.65 [0.47;0.91]** | **0.011** |
| CCL3 | CCL3 | 0.93 [0.64;1.35] | 0.706 | 0.82 [0.59;1.14] | 0.243 |
| **CCL5** | **CCL5** | 0.83 [0.55;1.23] | 0.346 | **0.61 [0.42;0.87]** | **0.007** |
| Chymase | CMA1 | 0.26 [0.02;4.34] | 0.347 | 0.37 [0.08;1.82] | 0.223 |
| **CXCL1** | **CXCL1** | **1.45 [1.04;2.02]** | **0.028** | 1.22 [0.82;1.82] | 0.319 |
| **CXCL10** | **CXCL10** | 1.06 [0.72;1.56] | 0.756 | **0.63 [0.43;0.93]** | **0.020** |
| CXCL2 | CXCL2 | 1.35 [0.96;1.88] | 0.083 | 1.01 [0.72;1.42] | 0.955 |
| **CXCL3** | **CXCL3** | **1.58 [1.16;2.16]** | **0.003** | 1.39 [0.96;2.02] | 0.080 |
| EPCR | PROCR | 1.05 [0.71;1.55] | 0.798 | 0.77 [0.56;1.06] | 0.114 |
| Heparin | SERPIND1 | 1.23 [0.90;1.67] | 0.193 | 1.13 [0.86;1.48] | 0.393 |
| **Histamine** | **HDC** | 0.58 [0.32;1.05] | 0.073 | **0.51 [0.30;0.86]** | **0.012** |
| IFN-G | IFNG | 1.34 [0.92;1.94] | 0.123 | 0.90 [0.55;1.48] | 0.683 |
| IL-1 | IL1A | 0.96 [0.66;1.40] | 0.820 | 1.00 [0.72;1.40] | 0.993 |
| **IL-13** | **IL13** | **1.39 [1.01;1.92]** | **0.042** | 1.30 [0.96;1.78] | 0.094 |
| **IL-16** | **IL16** | **0.65 [0.44;0.96]** | **0.031** | **0.56 [0.40;0.79]** | **0.001** |
| IL-17 | IL17A | 0.90 [0.61;1.35] | 0.622 | 0.73 [0.50;1.07] | 0.103 |
| **IL-4** | **IL4** | **1.47 [1.01;2.12]** | **0.042** | **1.47 [1.01;2.14]** | **0.046** |
| IL-5 | IL5 | 1.05 [0.75;1.47] | 0.777 | 1.00 [0.73;1.38] | 0.994 |
| IL-6 | IL6 | 1.35 [0.96;1.88] | 0.082 | 1.02 [0.67;1.56] | 0.909 |
| **IL-8** | **CXCL8** | **1.78 [1.23;2.57]** | **0.002** | **1.60 [1.14;2.25]** | **0.007** |
| **LTB** | **ALOX5** | 0.76 [0.52;1.11] | 0.155 | **0.70 [0.50;0.97]** | **0.033** |
| **OX40L** | **TNFSF4** | **1.42 [1.01;1.99]** | **0.041** | **1.48 [1.08;2.01]** | **0.014** |
| PGD2 | PTGDS | 0.78 [0.52;1.15] | 0.211 | 0.76 [0.54;1.06] | 0.107 |
| PGE2 | PTGES | 0.90 [0.60;1.35] | 0.606 | 0.76 [0.53;1.11] | 0.154 |
| PGI2 | PTGIS | 1.34 [0.94;1.92] | 0.103 | 1.11 [0.83;1.47] | 0.484 |
| **Substance P** | **PTGS2** | **1.52 [1.07;2.16]** | **0.020** | 1.31 [0.88;1.94] | 0.179 |
| TGF-B | TGFB1 | 0.99 [0.69;1.42] | 0.939 | 0.96 [0.70;1.32] | 0.808 |
| **TNF** | **TNF** | 0.85 [0.61;1.20] | 0.360 | **0.69 [0.49;0.97]** | **0.031** |
| **Tryptase** | **TPSAB1** | **0.63 [0.40;1.00]** | **0.048** | **0.54 [0.36;0.81]** | **0.003** |
| VEGF-A | VEGFA | 1.27 [0.86;1.87] | 0.230 | 1.22 [0.89;1.67] | 0.227 |
| VEGF-B | VEGFB | 0.99 [0.68;1.47] | 0.999 | 1.09 [0.78;1.51] | 0.621 |
| **VEGF-C** | **VEGFC** | **1.54 [1.00;2.35]** | **0.048** | 1.40 [0.98;2.01] | 0.064 |
| **VEGF-D** | **VEGFD** | **0.65 [0.45;0.95]** | **0.025** | **0.63 [0.45;0.89]** | **0.009** |

**Supplemental Figure 1.**
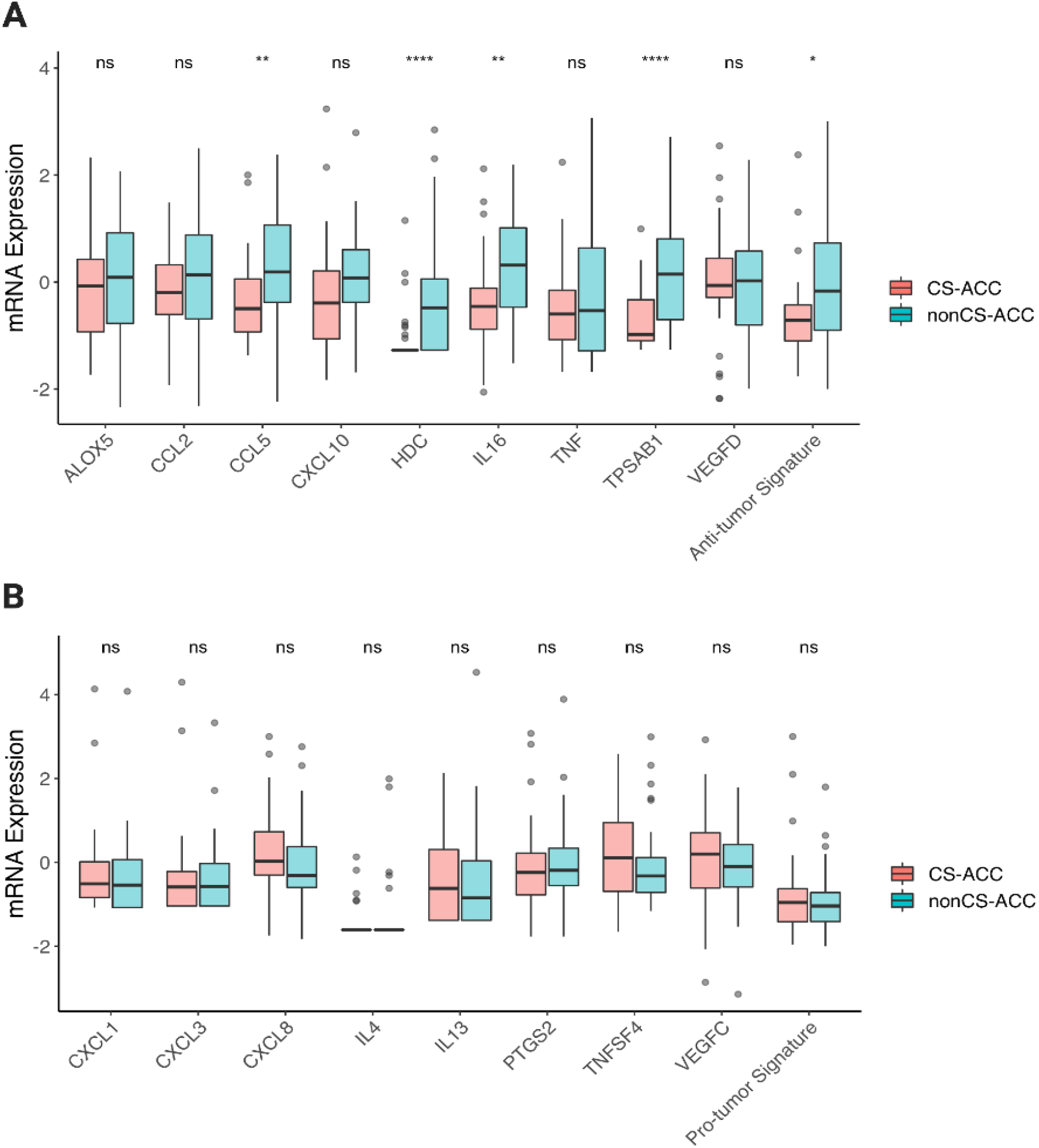
Differential Mast Cell Signaling Gene (MCSG) Expression by Cortisol Secretion in Adrenocortical Carcinoma. **A**. Anti-tumor MCSG mRNA expression in ACC tumors and stratified into subgroups by tumor cortisol secretion. **B**. Pro-tumor MCSG signaling genes mRNA expression in ACC tumors and stratified into subgroups by tumor cortisol secretion. ns = p value >0.05; * = p value <0.05, ** = p value <0.01, *** = p value <0.001, **** = p value <0.0001.

---

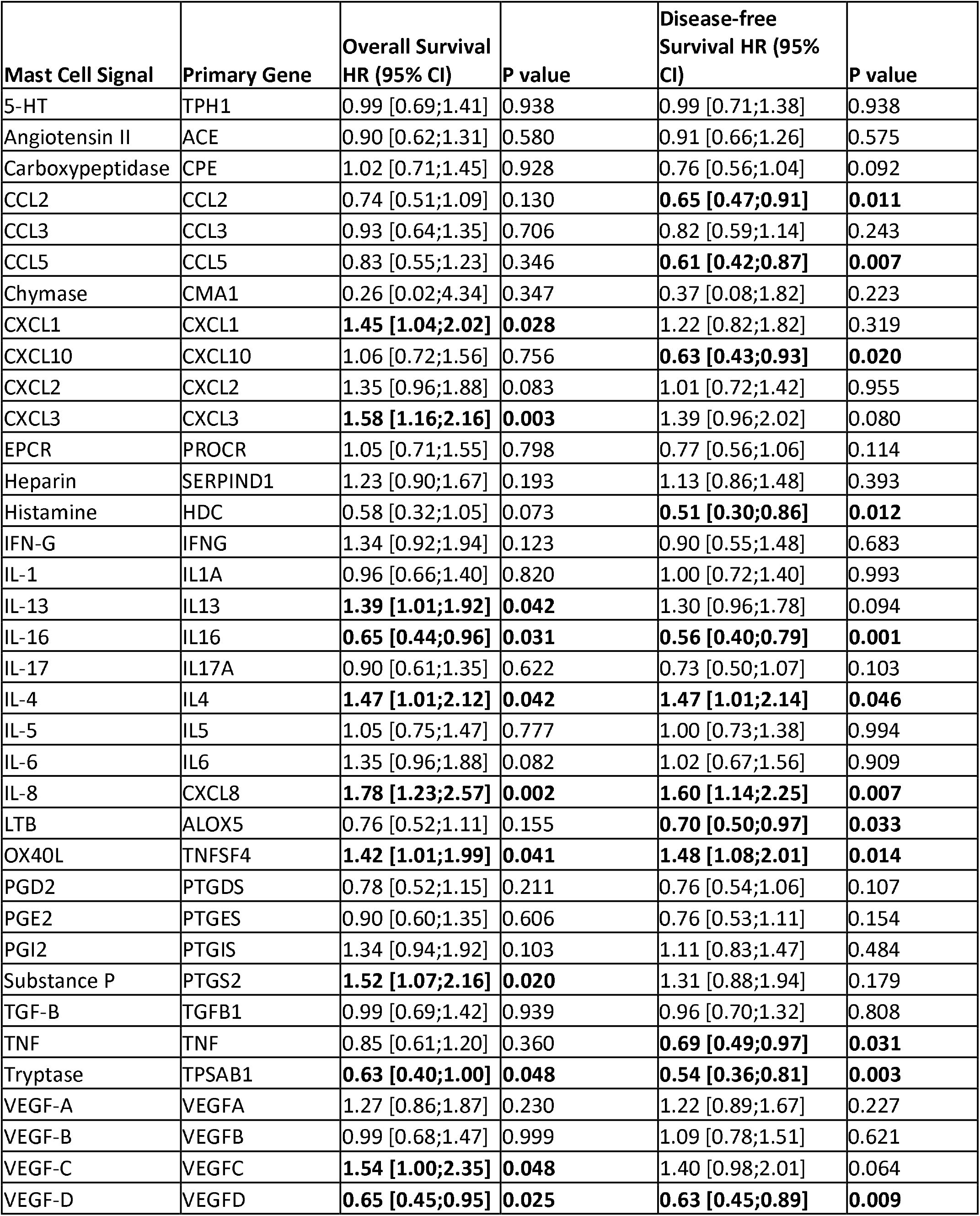

| Mast Cell Signal | Primary Gene | Overall Survival<br>HR (95% CI) | P value | Disease-free<br>Survival HR (95%<br>CI) | P value |
| --- | --- | --- | --- | --- | --- |
| 5-HT | TPH1 | 0.99 [0.69;1.41] | 0.938 | 0.99 [0.71;1.38] | 0.938 |
| Angiotensin II | ACE | 0.90 [0.62;1.31] | 0.580 | 0.91 [0.66;1.26] | 0.575 |
| Carboxypeptidase | CPE | 1.02 [0.71;1.45] | 0.928 | 0.76 [0.56;1.04] | 0.092 |
| CCL2 | CCL2 | 0.74 [0.51;1.09] | 0.130 | **0.65 [0.47;0.91]** | **0.011** |
| CCL3 | CCL3 | 0.93 [0.64;1.35] | 0.706 | 0.82 [0.59;1.14] | 0.243 |
| CCL5 | CCL5 | 0.83 [0.55;1.23] | 0.346 | **0.61 [0.42;0.87]** | **0.007** |
| Chymase | CMA1 | 0.26 [0.02;4.34] | 0.347 | 0.37 [0.08;1.82] | 0.223 |
| CXCL1 | CXCL1 | **1.45 [1.04;2.02]** | **0.028** | 1.22 [0.82;1.82] | 0.319 |
| CXCL10 | CXCL10 | 1.06 [0.72;1.56] | 0.756 | **0.63 [0.43;0.93]** | **0.020** |
| CXCL2 | CXCL2 | 1.35 [0.96;1.88] | 0.083 | 1.01 [0.72;1.42] | 0.955 |
| CXCL3 | CXCL3 | **1.58 [1.16;2.16]** | **0.003** | 1.39 [0.96;2.02] | 0.080 |
| EPCR | PROCR | 1.05 [0.71;1.55] | 0.798 | 0.77 [0.56;1.06] | 0.114 |
| Heparin | SERPIND1 | 1.23 [0.90;1.67] | 0.193 | 1.13 [0.86;1.48] | 0.393 |
| Histamine | HDC | 0.58 [0.32;1.05] | 0.073 | **0.51 [0.30;0.86]** | **0.012** |
| IFN-G | IFNG | 1.34 [0.92;1.94] | 0.123 | 0.90 [0.55;1.48] | 0.683 |
| IL-1 | IL1A | 0.96 [0.66;1.40] | 0.820 | 1.00 [0.72;1.40] | 0.993 |
| IL-13 | IL13 | **1.39 [1.01;1.92]** | **0.042** | 1.30 [0.96;1.78] | 0.094 |
| IL-16 | IL16 | **0.65 [0.44;0.96]** | **0.031** | **0.56 [0.40;0.79]** | **0.001** |
| IL-17 | IL17A | 0.90 [0.61;1.35] | 0.622 | 0.73 [0.50;1.07] | 0.103 |
| IL-4 | IL4 | **1.47 [1.01;2.12]** | **0.042** | **1.47 [1.01;2.14]** | **0.046** |
| IL-5 | IL5 | 1.05 [0.75;1.47] | 0.777 | 1.00 [0.73;1.38] | 0.994 |
| IL-6 | IL6 | 1.35 [0.96;1.88] | 0.082 | 1.02 [0.67;1.56] | 0.909 |
| IL-8 | CXCL8 | **1.78 [1.23;2.57]** | **0.002** | **1.60 [1.14;2.25]** | **0.007** |
| LTB | ALOX5 | 0.76 [0.52;1.11] | 0.155 | **0.70 [0.50;0.97]** | **0.033** |
| OX40L | TNFSF4 | **1.42 [1.01;1.99]** | **0.041** | **1.48 [1.08;2.01]** | **0.014** |
| PGD2 | PTGDS | 0.78 [0.52;1.15] | 0.211 | 0.76 [0.54;1.06] | 0.107 |
| PGE2 | PTGES | 0.90 [0.60;1.35] | 0.606 | 0.76 [0.53;1.11] | 0.154 |
| PGI2 | PTGIS | 1.34 [0.94;1.92] | 0.103 | 1.11 [0.83;1.47] | 0.484 |
| Substance P | PTGS2 | **1.52 [1.07;2.16]** | **0.020** | 1.31 [0.88;1.94] | 0.179 |
| TGF-B | TGFB1 | 0.99 [0.69;1.42] | 0.939 | 0.96 [0.70;1.32] | 0.808 |
| TNF | TNF | 0.85 [0.61;1.20] | 0.360 | **0.69 [0.49;0.97]** | **0.031** |
| Tryptase | TPSAB1 | **0.63 [0.40;1.00]** | **0.048** | **0.54 [0.36;0.81]** | **0.003** |
| VEGF-A | VEGFA | 1.27 [0.86;1.87] | 0.230 | 1.22 [0.89;1.67] | 0.227 |
| VEGF-B | VEGFB | 0.99 [0.68;1.47] | 0.999 | 1.09 [0.78;1.51] | 0.621 |
| VEGF-C | VEGFC | **1.54 [1.00;2.35]** | **0.048** | 1.40 [0.98;2.01] | 0.064 |
| VEGF-D | VEGFD | **0.65 [0.45;0.95]** | **0.025** | **0.63 [0.45;0.89]** | **0.009** |

|  |  | Low Mast Cells<br>N=64 | High Mast Cells<br>N=15 | p-value |
| --- | --- | --- | --- | --- |
| **Age (years) -Median [IQR]** |  | 48.0 [35.5;60.0] | 50.0 [33.5;53.0] | 0.871 |
| **Sex** | Female | 40 (62.5%) | 8 (53.3%) | 0.718 |
|  | Male | 24 (37.5%) | 7 (46.7%) |  |
| **Race** | White | 57 (89.1%) | 9 (60.0%) | 0.014 |
|  | Asian | 1 (1.56%) | 0 (0.0%) |  |
|  | Black | 0 (0.00%) | 1 (6.7%) |  |
|  | Unknown | 6 (9.4) | 5 (33.3%) |  |
| **Clinical Stage** | Stage I | 7 (10.9%) | 2 (13.3%) | 0.622 |
|  | Stage II | 28 (43.8%) | 9 (60.0%) |  |
|  | Stage III | 15 (23.4%) | 1 (6.7%) |  |
|  | Stage IV | 12 (18.8%) | 3 (20.0%) |  |
|  | Unknown | 2 (3.12%) | 0 (0.0%) |  |
| **T Stage** | T1 | 7 (10.9%) | 2 (13.3%) | 0.821 |
|  | T2 | 32 (50.0%) | 10 (66.7%) |  |
|  | T3 | 7 (10.9%) | 1 (6.7%) |  |
|  | T4 | 16 (25.0%) | 2 (13.3%) |  |
|  | Unknown | 2 (3.1%) | 0 (0.0%) |  |
| **N Stage** | N0 | 54 (84.4%) | 14 (93.3%) | 0.791 |
|  | N1 | 8 (12.5%) | 1 (6.7%) |  |
|  | Unknown | 8 (12.5%) | 0 (0.0%) |  |
| **M Stage** | M0 | 50 (78.1%) | 12 (80.0%) | 1.00 |
|  | M1 | 12 (18.8%) | 3 (20.0%) |  |
|  | Unknown | 2 (3.12%) | 0 (0.0%) |  |
| **Excess Hormone Secretion** | Cortisol | 16 (25.4%) | 0 (0.0%) | 0.047 |
|  | Androgen + Cortisol | 13 (20.6%) | 3 (20.0%) |  |
|  | Androgen | 8 (12.7%) | 0 (0.0%) |  |
|  | Estrogen | 1 (1.6%) | 1 (6.7%) |  |
|  | Mineralocorticoids | 2 (3.2%) | 1 (6.7%) |  |
|  | Mineralocorticoid + Cortisol | 1 (1.6%) | 0 (0.0%) |  |
|  | None | 21 (33.3%) | 9 (60.0%) |  |
| **Excess Cortisol (including mixed) Secretion** | No | 34 (53.1%) | 12 (80%) | 0.108 |
|  | Yes | 30 (46.9%) | 3 (20.0%) |  |
| **Laterality** | Left | 38 (59.4%) | 7 (46.7%) | 0.545 |
|  | Right | 26 (40.6%) | 8 (53.3%) |  |
| **Fraction of Genome Altered -Median [IQR]** |  | 0.6 [0.4;0.9] | 0.42 [0.3;0.8] | 0.276 |
| **Mutation Count -Median [IQR]** |  | 87.0 [72.5;118] | 81.5 [57.5;92.8] | 0.222 |
| **Mitoses Count /mm<sup>3</sup> -Median [IQR]** |  | 7.00 [5.0;23.0] | 7.00 [4.0;15.0] | 0.556 |
| **Mitotic Rate** | < 5/50 HPF | 24 (37.5%) | 5 (33.3%) | 0.777 |
|  | > 5/50 HPF | 32 (50.0%) | 9 (60.0%) |  |
|  | Unknown | 8 (12.5%) | 1 (6.8%) |  |
| **Tumor Necrosis** | Absent | 14 (21.9%) | 3 (20.0%) | 1.00 |
|  | Present | 46 (71.9%) | 11 (73.3%) |  |
| **Weiss Score -Median [IQR]** |  | 5.50 [3.8;7.3] | 7.00 [5.0;7.0] | 0.545 |
| **Venous Invasion** | Absent | 31 (48.4%) | 8 (53.3%) | 0.849 |
|  | Present | 26 (40.6%) | 5 (33.3%) |  |
|  | Unknown | 7 (10.9%) | 2 (13.3%) |  |
| **Neoadjuvant Therapy** | No | 63 (98.4%) | 1 (1.6%) | 1.000 |
|  | Yes | 1 (1.56%) | 0 (0.0%) |  |
| **Resection Margin** | R0 | 43 (67.2%) | 12 (80.0%) | 0.866 |
|  | R1 | 5 (7.8%) | 1 (6.7%) |  |
|  | R2 | 7 (10.9%) | 2 (13.3%) |  |
|  | RX | 6 (9.4%) | 0 (0.0%) |  |
|  | Unknown | 3 (4.7%) | 0 (0.0%) |  |
| **Adjuvant Therapy** | No | 20 (31.2%) | 5 (33.3%) | 0.662 |
|  | Yes | 42 (65.6%) | 9 (60.0%) |  |
|  | Unknown | 2 (3.12%) | 1 (6.7%) |  |
| **Mitotane Therapy** | No | 21 (32.8%) | 5 (33.3%) | 0.231 |
|  | Yes | 41 (64.1%) | 8 (53.3%) |  |
|  | Unknown | 2 (3.12%) | 2 (13.3%) |  |
| **Radiation Therapy** | No | 47 (73.4%) | 12 (80.0%) | 0.866 |
|  | Yes | 14 (21.9%) | 1 (6.7%) |  |
|  | Unknown | 3 (4.7%) | 2 (13.3%) |  |
| **New Neoplasm Following Initial Treatment** | No | 26 (44.1%) | 12 (85.7%) | 0.012 |
|  | Yes | 33 (55.9%) | 2 (14.3%) |  |

